# TP63-TRIM29 axis regulate enhancer methylation and chromosomal instability in prostate cancer

**DOI:** 10.1101/2022.03.08.483320

**Authors:** R. Sultanov, A. Mulyukina, O. Zubkova, A. Fedoseeva, A. Bogomazova, K. Klimina, A. Larin, T. Zatsepin, T. Prikazchikova, E. Sharova, E. Generozov, V. Govorun, M. Lagarkova, G. Arapidi

**Affiliations:** Center for Precision Genome Editing and Genetic Technologies for Biomedicine, Federal Research and Clinical Center of Physical-Chemical Medicine of Federal Medical Biological Agency, Moscow, Russia; Federal Research and Clinical Center of Physical-Chemical Medicine of Federal Medical Biological Agency, Moscow, Russia; Shemyakin-Ovchinnikov Institute of Bioorganic Chemistry of the Russian Academy of Sciences, Moscow, Russia; Moscow Institute of Physics and Technology (National Research University), Dolgoprudny, Russia; Skolkovo Institute of Science and Technology (National Research University), Moscow, Russia; Department of Chemistry, Lomonosov Moscow State University, Moscow, Russia

**Keywords:** TP63, TRIM29, epigenetic regulation, prostate cancer, WGCNA

## Abstract

Prostate adenocarcinoma (PRAD) is the second most common cause of cancer-related deaths in men. PRAD is often characterized by DNA methylation variability and a high rate of large genomic rearrangements. To elucidate the reasons behind such high variance, we used weighted gene co-expression network analysis for integration RNA-seq, DNA methylation and copy number alterations data from The Cancer Genome Atlas PRAD. Our results show that only a single cluster of co-expressed genes is associated with genomic and epigenomic instability. Within this cluster, TP63 and TRIM29 are key transcription regulators. We revealed that TP63 regulates the level of enhancer methylation in prostate basal epithelium cells. TRIM29 forms a complex with TP63 and together regulate the expression of genes specific to the prostate basal epithelium. Moreover, TRIM29 binds DNA repair proteins and prevents formation of the *TMPRSS2:ERG* gene fusion typically observed in PRAD. Therefore, the study shows that TRIM29 and TP63 are important regulators maintaining the identity of the basal epithelium under physiological conditions. Finally, we uncover the role of TRIM29 in PRAD development.

## Introduction

Prostate adenocarcinoma (PRAD) is the most frequently diagnosed cancer in men and the second leading cause of cancer mortality affecting men worldwide (Sung et al. 2021). In contrast to other types of cancer, PRAD is characterized by a low number of recurrent single nucleotide variations (SNVs). At the same time, it exhibits high genomic and chromosomal instability accompanied by a high frequency of gene fusions (e.g., *TMPRSS2:ERG* is found in ∼50% of patients) and high epigenomic variability (Kluth et al. 2013; Taylor et al. 2010; Cancer Genome Atlas Research Network 2015). Many studies have demonstrated that epigenetic changes, including DNA methylation and histone modifications, are required for the initiation and progression of PRAD (Kgatle et al. 2016; Ruggero et al. 2018; Ferguson et al. 2015). Methylome changes are often caused by somatic mutations in genes of the DNA methylation-demethylation system and chromatin modulator system (*DNMT3B*, *TET2*, *BRAF*, *IDH1*) (Zhao et al. 2020). Altering the activity of transcription factors such as androgen receptor (AR) or ETS-related gene (ERG) can also lead to dramatic changes in the epigenetic landscape (Stelloo et al. 2018). Interestingly, changes in methylation patterns may also affect the activity of genes involved in cell cycle control, response to hormones, and DNA damage repair (Börno et al. 2012; Kobayashi et al. 2011; Zhao et al. 2020). Chromosomal instability in PRAD is often the result of a faulty DNA damage repair system, leading to amplification and deletion of large loci and chromosome arms (Stopsack et al. 2019; Cancer Genome Atlas Research Network 2015; Taylor et al. 2010). PRAD exhibits amplification of such oncogenes as MYC and AR and the formation of the *TMPRSS2:ERG* gene fusion, which is the most common recurrent somatic mutation in PRAD. *TMPRSS2:ERG* gene fusion has pioneering transcription factor properties that can alter the epigenetic landscape and cause (Babu and Fullwood, 2017; Kron et al., 2017), MYC and AR overexpression (Guo et al. 2021; Stelloo et al. 2018). Thus, chromosomal instability and changes in DNA methylation can mutually influence each other. However, molecular mechanisms that lead to high chromosomal instability and epigenomic variability remain unclear. In recent years, continued efforts from several projects like The Cancer Genome Atlas (TCGA), International Cancer Genome Consortium (ICGC), Chinese Prostate Cancer Genome and Epigenome Atlas (CPGEA), and Taylor dataset (Taylor et al. 2010) has helped gathering multiple PRAD-related omics data. Comprehensive profiling of RNA seq, DNA methylation, and somatic mutations data from these projects has revealed seven distinct subtypes of PRAD (Cancer Genome Atlas Research Network 2015). However, integrating various omics data and interpreting the results still presents some challenges. RNA profiling is one of the significant genomic tools to study molecular mechanisms underlying various diseases. RNA-seq captures the average gene expression profile from the tissue. However, due to the highly heterogeneous nature of PRAD tumors, the resulting average gene expression profile from tumor tissues is not always a direct depiction of their signature molecular events. This issue can be addressed by searching for clusters (or modules) of co-expressed genes to be further associated with clinical data and phenotypes. Weighted gene co-expression network analysis (WGCNA) (Langfelder and Horvath 2008) allows assembly of thousands of genes with similar patterns of expression into clusters. These clusters can be associated with various phenotypes or other omics data for correlative or quantitative studies. The WGCNA approach has already been used to search for important gene hubs, biomarkers, therapeutic targets, or integration with omics data for PRAD (Hou et al. 2021; Ohandjo et al. 2019; S. Li et al. 2017). In addition, the WGCNA algorithm has been proven to be an exemplary method for efficient generation of hypotheses (Hou et al. 2021; Ohandjo et al. 2019; S. Li et al. 2017).

Here, we used the WGCNA correlation network analysis algorithm to search for clusters of co-expressed genes associated with epigenetic variability and chromosomal instability in PRAD using the TCGA PRAD data set. Further analysis of the module using GO, TF-enrichment analysis, and KEGG analysis identified the TP63 cluster responsible for regulating the cluster of genes associated with epigenetic variability and genome instability in PRAD. We showed that TRIM29 interacts with TP63 and regulates the expression of the TP63 cluster, which is associated with variability in DNA methylation and chromosomal instability. Using *TP63* and *TRIM29* gene knockdown and overexpression in two cell lines of the normal basal epithelium of the prostate and two cell lines of PRAD we showed that TP63 directly regulates methylation levels of CpG sites in enhancers specific for the basal epithelium of the prostate. In contrast, TRIM29 regulates the activity of transcription factor TP63. Furthermore, we show that in the TP63 cluster, TRIM29 performs the genome protection functions against chromosomal instability.

## Results

### TP63 regulates the gene cluster associated with epigenetic variability and chromosomal instability in PRAD

To unveil a cluster of co-expressed genes associated with epigenomic variability and chromosomal instability, we analyzed RNA-seq, DNA methylation (Infinium MethylationBeadChip450K), and copy number variation (CNV, Affymetrix 6.0) datasets of The Cancer Genome Atlas (TCGA) for prostate adenocarcinoma (PRAD). In brief, we determined 13 co-expressed gene clusters in the RNA-seq data using the WGCNA package (Langfelder and Horvath 2008). To determine association of clusters with differentially expressed genes between tumor and adjacent non-tumor tissues, we analyzed changes in methylation of CpG sites and chromosomal instability index (see Methods for details). Based on the established criteria, subsequent analysis yielded a single cluster of 143 genes that were found to be heavily correlated with 1,645 differentially methylated CpG sites (Figure 1; Table S1, S2).

**Figure 1.**
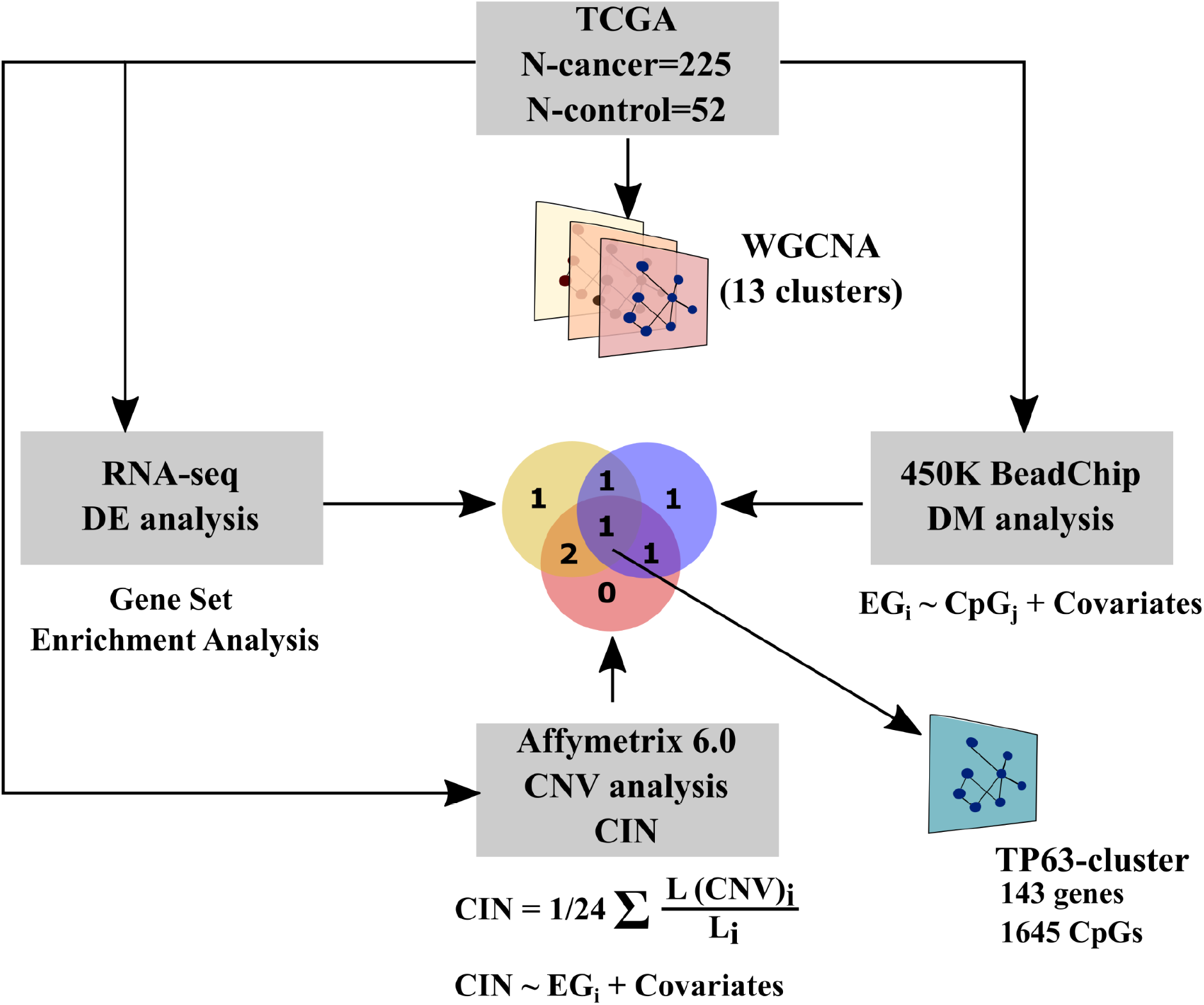
Analysis of clusters of co-expressed genes associated with epigenetic variability and chromosomal instability in PRAD. Briefly, 13 clusters of co-expression genes were generated from TCGA PRAD datasets using WGCNA. To identify clusters associated with epigenetic variability and chromosomal instability, GSEA, regression analysis between eigengene of cluster and CpG-site methylation or chromosomal instability index was performed. DE - differential expression, DM - differential methylation, CIN - chromosomal instability index.

Based on Gene set enrichment analysis (GSEA) (Subramanian et al. 2005), the group of genes downregulated in PRAD was significantly enriched with genes of this cluster (Figure S1A). Out of 1,645 CpG sites associated with the cluster, 1539 sites (94%) were hypermethylated. GO term analysis revealed genes that were enriched for the “epidermis development,” “epithelial cell development,” and “focal adhesion processes” (Table S2; Figures 2A, B).

**Figure 2.**
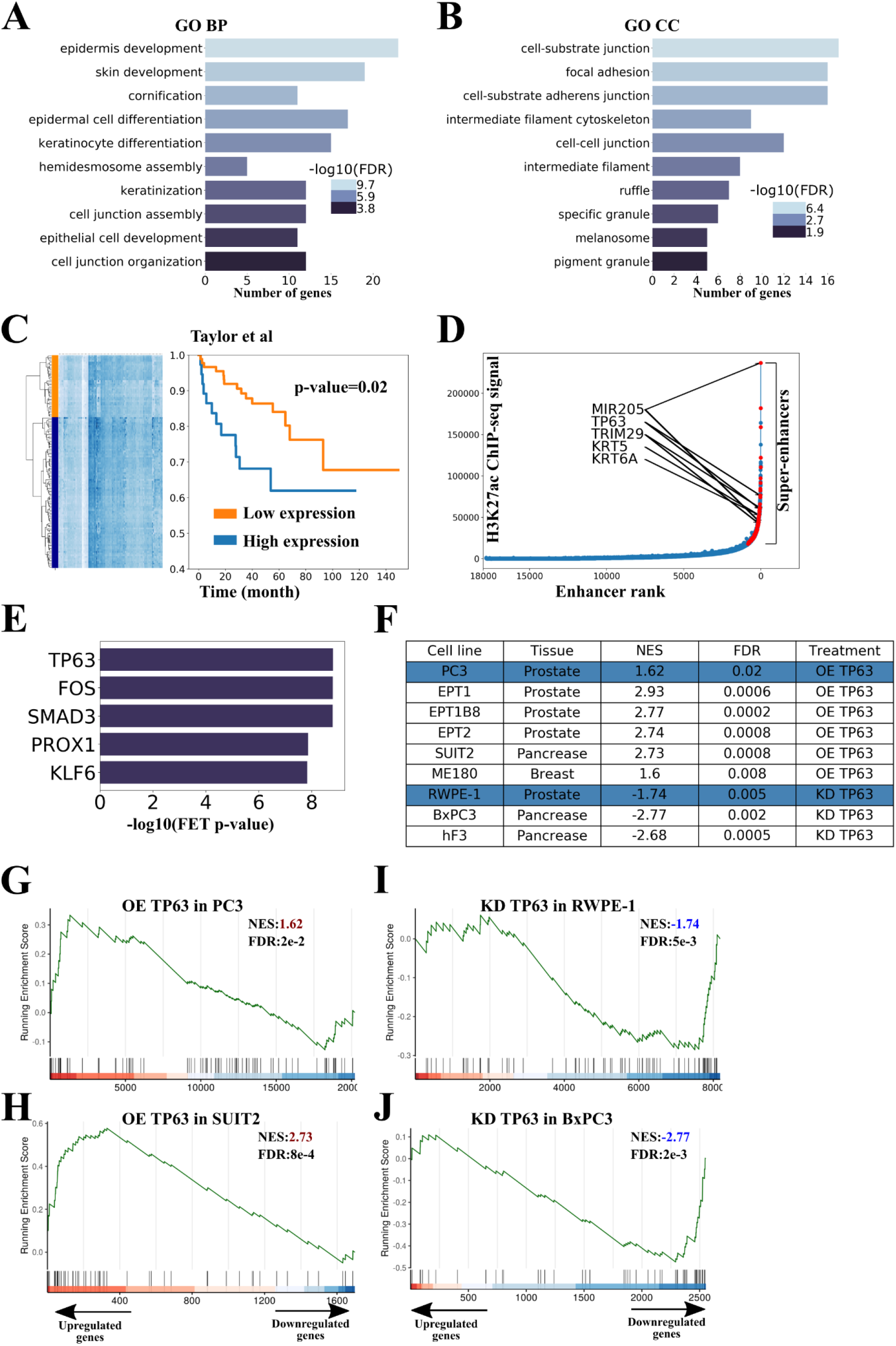
TP63 regulates the cluster of genes associated with epigenetic variability and genome instability in PRAD. 2A, B. Gene Ontology (GO) analysis of the cluster genes. GO terms related to biological process (A), and cellular components (B) are shown. FDR is indicated with a color scale. 2C. Biochemical recurrence-free survival curves of patients from the Taylor dataset <u>(Taylor et al.</u> <u>2010)</u> stratified by expression of TP63-cluster genes. P-value was calculated using the log-rank test. 2D. Hockey stick plot based on input-normalized H3K27ac signals in PrEC cell line. Super-enhancer-associated cluster genes are highlighted with red. 2E. Results of TF-enrichment analysis (CHEA3) for the cluster of interest. FET - Fisher’s exact test. 2F. Table summarizing RNA-seq data evaluating the cluster genes signature upon *TP63* overexpression (OE) or knockdown (KD) in the indicated cell lines. Blue rows indicate data from this study. 2G-H. Gene set enrichment analysis (GSEA) plots evaluating the cluster genes signature upon *TP63* overexpression (OE) in PC3 (PRAD, 2G) and SUIT2 (GSE115462, pancreatic cancer, 2H). NES - Normalized enrichment scores. 2I-J. GSEA plots evaluating the cluster genes signature upon *TP63* knockdown (KD) in RWPE-1 (prostate basal epithelium, 2I) and BxPC3 (GSE115462, pancreatic cancer, 2J).

Similarly, using the independent PRAD RNA-seq dataset obtained by Taylor and co-authors, we showed that high expression of the cluster genes is associated with the late biochemical recurrence. In contrast, low expression of the cluster genes is associated with the earlier biochemical recurrence (Figure 2C) (Taylor et al. 2010). When compared with prostate scRNA-seq data (Henry et al. 2018), we showed that the cluster was significantly enriched with signatures of basal prostate epithelium, but not luminal epithelium or stromal cells (Fisher exact test OR = 50, p-value = 3e-11). Finally, we also confirmed that the cluster is regulated by super-enhancers (SEs) specific for the prostate basal epithelium (43 genes, OR=3, p-value=1.5e-5, Figure 2D), which may indicate that the cluster is associated with cell lineage specification and cell fate decision functions (Hnisz, Day, and Young 2016).

Often, clusters of co-expressed genes are regulated by only one or few transcription factors. We used the CHEA3 program (Keenan et al. 2019) and identified the top five most likely candidates regulating the cluster: TP63, FOS, SMAD3, PROX1, KLF6 (Figure 2E). Since only TP63 belongs to the cluster of co-expressed genes, we hypothesized that it could serve as a probable regulator of the cluster, as TP63 is a recognized master regulator of epithelium development (Somerville et al. 2018; Y.-Y. Jiang et al. 2020; Y. Jiang et al. 2018; Kouwenhoven et al. 2015). Additionally, expression of TP63 is considerably decreased in PRAD and absent in all studied PRAD cell lines (Figure S1B and S1C). To uncover the effects of TP63 on the cluster, we overexpressed *TP63* in the PC3 PRAD cell line (similar to most PRAD cell lines, PC3 cells lack endogenous expression of *TP63*) and knocked down *TP63* in RWPE-1 normal prostate epithelial cells (Figure S1D) and performed RNA-seq analysis. Expression of *TP63* led to a considerable increase in the expression of the cluster genes in PC3 cells (Figures 2F and 2G). On the contrary, knockdown of *TP63* resulted in a significant decrease in the cluster gene expression in RWPE-1 cells (Figures 2F and 2H). We proposed that the association between *TP63* and the cluster gene expression could be common across different cancer types. Through this process, we demonstrated that expression of the cluster genes depended on *TP63* in pancreatic cancer (SUIT2, BxPC3, hF3; Figures 2F-H, Table S1), prostate cancer (EPT1, EPT1B8, EPT2; Figure 2F, 2I-J, Table S1), and cervical cancer (ME180; Figure 2F, Table S1) cell lines. To show that *TP63* directly regulates the cluster genes, we performed ChIP-seq with anti-TP63 antibodies using RWPE-1 normal prostate basal epithelial cell line and PC3 PRAD cell line with overexpression of *TP63* (PC3-TP63). In both normal basal epithelium and PC3-TP63 cells, the cluster genes were significantly enriched with ChIP-seq peaks of TP63 (Fisher exact test OR=5.6 and OR=5.7, p-value=6.3e-17 and p-value=7.8e-21, respectively). Therefore, we confirmed that transcription of the cluster (further referred as TP63 cluster) is regulated by TP63.

### CpG sites associated with the TP63 cluster belong to TP63-dependent enhancers and super-enhancers

Here we proposed that transcription of TP63 cluster is associated with changes in methylation of 1645 CpG sites (TP63 CpG sites from now on, Table S3). Based on this assumption, we performed GO analysis of genes located in proximity to these CpG sites. Our subsequent analysis confirmed their association with prostate development, morphogenesis, and cancer (Figures 3A, 3B, Table S3). To determine the functional role of these sites, we used open ChIP-seq data on histone modifications H3K27ac, H3K4me1, and H3K4me3 for normal prostate basal epithelium (RWPE1, PrEC; Table S1) and PRAD (LNCaP, PC3; Table S1) cell lines. TP63 CpG sites were found to be enriched with enhancer features in PrEC and RWPE-1 cell lines, but not in PRAD cells. They were localized mainly in distal (non-promoter) regions compared to random CpG sites (Figures 3C, 3D). Using the activity-by-contact (ABC) models (Fulco et al. 2019), we predicted enhancers for the PrEC normal basal epithelium cell line. Out of 1645 TP63 CpG sites associated with the cluster, 175 CpG sites resided in the ABC enhancer region (Fisher exact test OR=3.8, p-value=2e-45). Moreover, TP63 CpG sites were considerably enriched with SEs of the PrEC cells line (Fisher exact test OR=5.1, p-value=2e-63). Therefore, we found that TP63 CpG sites belonged to enhancers specific for prostate basal epithelium. A survey for transcription factor motifs within 100 bp from TP63 CpG sites showed significant enrichment of motifs for the binding of the p53 family of transcription factors (Figure 3E). In addition, TP63 CpG sites showed intensive TP63 ChIP-seq signals in the RWPE-1 and PC3-TP63 cell lines (Figure 3F). Therefore, this data confirmed that TP63 CpG sites are located in TP63-dependent enhancers of basal epithelium.

**Figure 3.**
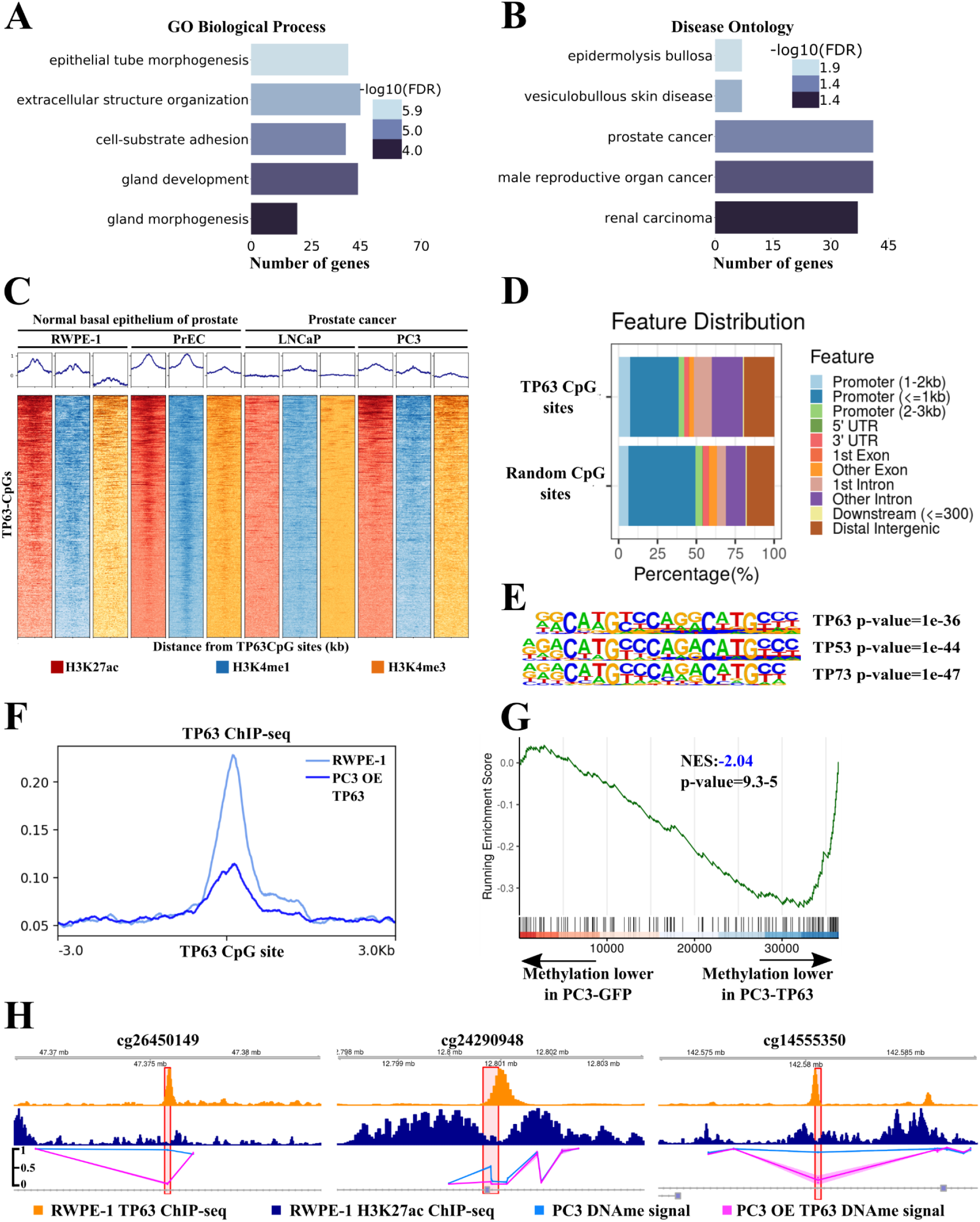
CpG sites associated with TP63 cluster lie in TP63-dependent enhancers and super-enhancers. 3A,B. Gene Ontology (GO) analysis of genes located close to TP63 CpG sites. GO terms related to biological process (A) and disease ontology (B) are shown. FDR is indicated with a color scale. 3C. ChIP-seq density plots of H3K27ac, H3K4me1, and H3K4me3 enrichments at TP63 CpG sites in the RWPE-1, PrEC, LNCaP, and PC3 cell lines. Each column represents a 6-kb interval centered on a CpG site. 3D. Bar-plot of genomic distribution of the TP63 CpG sites (top) and random CpG sites (bottom). 3E. Representation of p53 protein family motifs enriched at TP63 CpG sites using HOMER. 3F. Mean TP63 ChIP-seq signal at TP63 CpG sites in the RWPE-1 cell line and PC3 with overexpression (OE) of *TP63*. 3G. GSEA-like plot for methylation data evaluating the TP63 CpG sites signature upon *TP63* overexpression in PC3. 3H. ChIP-seq profiles of H3K27ac (top track) and TP63 (middle track) in RWPE-1 cell line and DNA methylation profile in PC3 cell line (bottom track) following overexpression (OE) of *TP63* or GFP as a control at three CpG sites.

Having established a link between TP63 and TP63 CpG sites, we evaluated the functional role of TP63 in regulating methylation of these CpG sites next. We performed whole-genome methylation analysis using the Illumina MethylationEPIC chip in the PC3 cell line with *TP63* overexpression. TP63 CpG sites were almost completely methylated in PC3 (Figure S2A). Upon induced expression of *TP63* in PC3, the level of TP63 CpG sites methylation significantly decreased (Figure 3G). CpG sites with decreased methylation level under TP63 overexpression belonged to TP63 ChIP-seq peaks in RWPE-1 and PC3-TP63 cell line (Figure 3H). To show that a decrease in the expression of *TP63* can lead to increased methylation of TP63 CpG sites, we chose three CpG sites that alter the levels of methylation in overexpressed *TP63* in PC3 and belong to TP63 ChIP-seq peaks in the RWPE-1 cell lines. We performed knockdown of the *TP63* gene in the RWPE-1 cell line and used MSRE-qPCR to show that methylation significantly increased in two of the three CpG sites (Figure S2B and S2C). Thus, both increase and decrease of TP63 expression can lead to changes in methylation of TP63 CpG sites. Collectively, these results strongly indicate that CpG sites associated with the TP63 cluster belong to TP63-dependent enhancers of prostate basal epithelium and their methylation level depends on TP63 expression.

### TRIM29 interacts with TP63 and regulates the expression of the TP63 cluster

Oncogenic or tumor suppressor properties of TP63 often depend on its protein partners in the cell (Yi et al. 2020). In squamous cell carcinoma (SCC), TP63 is an oncogene, and interacts with pioneer transcription factors as SOX2 and KLF5 (Y.-Y. Jiang et al. 2020; Y. Jiang et al. 2018). By contrast, TP63 is a tumor suppressor in PRAD and cervical cancer (Zhou et al. 2020). Since the function of TP63 partners still remain relatively unknown in these types of cancer, we decided to identify potential TP63 co-regulators in the TP63 cluster. According to Langfelder and Horvath, the eigengene of the cluster of co-expressed genes (pattern of cluster expression) has the highest correlation with the expression rate of potential regulators of the co-expressed gene cluster in WGCNA (Langfelder and Horvath 2008). *KRT5* and *TRIM29* are the top two genes in terms of degree of expression correlation with TP63 cluster eigengene expression (Figure 4A). KRT5 is a cytokeratin marker of basal epithelium. TRIM29 is a ubiquitin ligase of the TRIM family which is a known partner of TP53 (a TP63 homologue) and regulates its transcription factor activity (Yuan et al. 2010). Therefore, TRIM29 can be an important regulator of the TP63 cluster. Analysis of four PRAD RNA-seq datasets (Taylor et al. 2010; Fraser et al. 2017; Cancer Genome Atlas Research Network 2015; Zhang et al. 2019) (Table S1) showed a strong correlation between *TRIM29* and *TP63* (Figure 4B). This observation concurs with TRIM29 regulating the basal invasive program *via* TP63 in bladder cancer (Palmbos et al. 2019). Thus, we aimed to decipher the effect TRIM29 has on the TP63 cluster in PRAD-related cell lines. To this end, we performed *TRIM29* knockdown in a prostate basal epithelium cell line RWPE-1 and induced expression of *TRIM29* in PC3 cell line followed by RNA-seq analysis. *TRIM29* knockdown in RWPE-1 led to a significant decrease in the expression of the TP63 cluster genes (Figure 4C), which agrees with previous observations from the dataset on *TRIM29* knockdown in normal breast epithelial cell lines HMEC and MCF10A (Table S1, Figure 4C, and Figure S3A). Interestingly, knockdown of *TP63* and *TRIM29* in RWPE-1 resulted in changes in the expression of overlapping gene sets of the cluster (Figure 4C) involved in processes such as epidermis development and differentiation, keratinocyte differentiation, and cell junction organization (Table S4). On the other hand, overexpression of *TRIM29* in PC3 did not affect the cluster gene expression. Yet, combined overexpression of *TP63* and *TRIM29* increased the number of differentially expressed genes of the TP63 cluster (Figure 4D).

**Figure 4.**
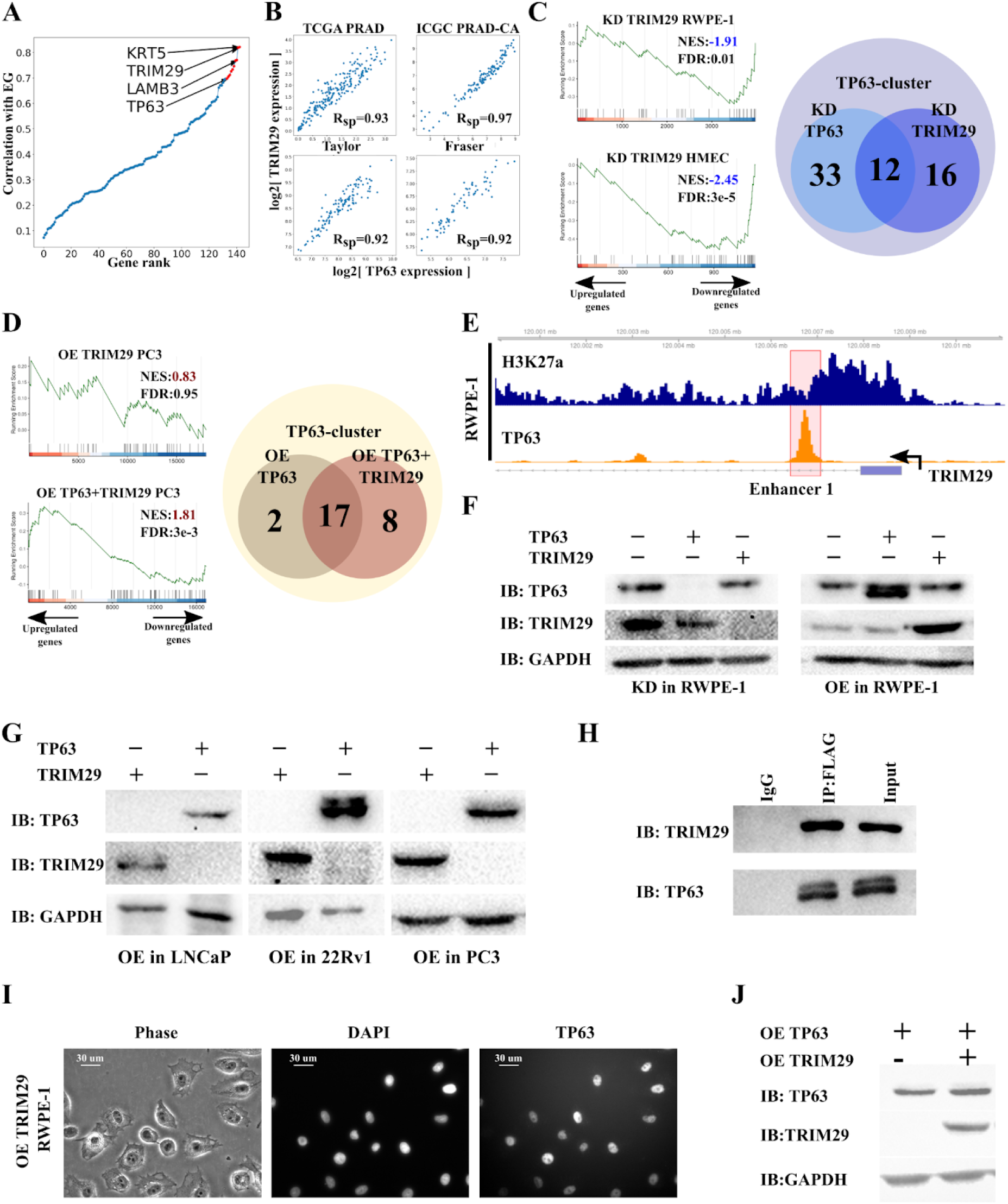
TRIM29 interacts with TP63 and regulates the expression of the TP63 cluster. 4A. Eigengene of TP63 cluster significantly correlates with the expression of *TRIM29*. 4B. *TP63* expression significantly correlates with the expression of *TRIM29* in four PRAD data sets. 4C. Comparison of *TP63* and *TRIM29* knockdowns (KDs). GSEA plots evaluating the TP63 cluster signature upon *TRIM29* knockdown in RWPE-1 (prostate basal epithelium, left-top) and HMEC (GSE71375, basal breast epithelium, left-bottom). Venn diagram showing the overlap of differentially expressed TP63 cluster genes upon *TP63* and *TRIM29* knockdown in the RWPE-1 cells (right). NES - Normalized enrichment scores. 4D. Comparison of *TP63* and *TP63*+*TRIM29* overexpression (OEs). GSEA plots evaluating the TP63 cluster signature upon *TRIM29* overexpression in PC3 (PRAD, left-top) and *TP63* and *TRIM29* simultaneous overexpression in PC3 (PRAD, left-bottom). Venn diagram showing the overlap of differentially expressed TP63 cluster genes upon *TP63* overexpression and *TP63* and *TRIM29* simultaneous overexpression in PC3 cells (right). 4E. ChIP-seq profiles of H3K27ac (top track) and TP63 (bottom track) in the RWPE-1 cell line of the *TRIM29* gene. Enhancer 1 of *TRIM29* in a red polygon. 4F. Immunoblot (IB) analysis of knockdown (KD) of *TP63* and *TRIM29* (left) and *TP63* and *TRIM29* overexpression (OE, right) in the RWPE-1 cells. 4G. Immunoblot (IB) analysis of overexpression (OE) of *TP63* and *TRIM29* in PRAD cell lines: LNCaP, 22Rv1, and PC3. 4H. Interaction between TRIM29 and TP63 was detected by co-immunoprecipitation (co-IP) in the PC3 cells overexpressing *TRIM29-FLAG* and *TP63*. 4I. Overexpression (OE) of *TRIM29* does not lead to the relocation of TP63 from the nucleus to the cytoplasm. Immunocytochemistry staining of TP63 in the RWPE-1 cells overexpressing *TRIM29*. 4J. Immunoblot (IB) analysis of overexpression (OE) of *TP63* and *TRIM29* simultaneously in PC3 cells.

Since TP63 and TRIM29 regulate the TP63 cluster simultaneously, we inquired whether TP63 and TRIM29 can mutually regulate each other. We proposed three possible ways of mutual regulation of the expression of these genes. 1) TP63 or TRIM29 can directly regulate the expression of each other. 2) TRIM29 could change the localization of TP63 on the cell, and 3) TRIM29 could act as ubiquitin-ligase, leading to degradation of TP63. Even though TP63 binds with TRIM29 enhancer in prostate basal epithelium and upon *TP63* overexpression in PC3 (Figure 4E, Figure S3B), knockdown of *TP63* in PRAD cell lines did not lead to an increased TRIM29 expression (Figures 4G–J). Overexpression or knockdown of *TRIM29* did not change the expression of TP63 (Figures 4G–J). Thus, in PRAD, TP63 and TRIM29 did not regulate each other at the transcription level.

We then postulated if TRIM29 can interact with TP63 and thus regulate its transcription factor activity. We induced overexpression of TRIM29-FLAG and TP63 in a PRAD cell line PC3 followed by immunoprecipitation with FLAG. Results showed that TP63 and TRIM29 interact with each other (Figure 4H). Furthermore, the mass spectrometry analysis of TRIM29 partner proteins confirmed the interaction between TRIM29 and TP63 in RWPE-1 (Table S5). Previous studies have shown that TRIM29 can lead to proteasomal degradation or change of localization of transcription factors (Q. Li et al. 2018; Cao et al. 2019; Xing et al. 2016). TRIM29 can translocate TP53 - homologue of TP63 from the nucleus to cytoplasm (Yuan et al. 2010). We hypothesize that TRIM29 might regulate TP63 activity by a similar mechanism. Hence, overexpression of TRIM29 in RWPE-1 did not lead to translocation of TP63 from the nucleus to cytoplasm (Figure 4I). Also, overexpression of *TRIM29* did not lead to degradation of TP63 (Figures 4F and 4J), as in the case of the STING protein (Q. Li et al. 2018). Consequently, TRIM29 does not regulate TP63 cluster expression by modulating TP63 abundance in the nucleus. All these data indicate that TRIM29 can modulate TP63 transcription factor activity and thus regulate cluster expression. It remains unclear, though, how the TP63 activity is regulated. Further research will need to investigate to determine how TP63 is regulated.

### Restoration of TRIM29 expression promotes the decrease of chromosomal instability in PRAD

Earlier, we demonstrate that the TP63 cluster is associated with increased chromosomal instability index and is not enriched with DNA repair proteins family of genes. We investigated two datasets that contained whole-genome screening of 28 genotoxic agents in retinal pigment epithelium (Olivieri et al. 2020) and two agents in HeLa cells (Schleicher et al. 2020). Interestingly, there was no significant association between the TP63 cluster and the response to genotoxic stress (Fisher exact test, Table S5), which led us to hypothesize that the association between TP63 cluster with chromosomal instability can be a consequence of the association of a cluster’s hub gene with chromosomal instability. Upon literature search, as expected, the most probable candidate for chromosomal instability regulator was TRIM29. Previously, TRIM29 was found to be a scaffold protein of DNA double-strand break repair system in HeLa (cervical cancer) (Masuda et al. 2015). Also, TRIM29 exhibited radioprotective function in a number of studies (Wang et al. 2014; Masuda et al. 2015; Yuan et al. 2010). Moreover, TRIM29 is a regulator of the TP63 cluster. Considering this, we decided to examine the effect of TRIM29 on chromosomal instability in PRAD. We first performed immunoprecipitation of TRIM29 from the nuclear fraction of RWPE-1 followed by Mass spectrometry analysis of the precipitate. Data showed that TRIM29 binds to DNA repair proteins that belong to mismatch excision repair, nucleotide excision repair, and homologous recombination families, as well as TP53BP1 (Table S5). Therefore, a low abundance of TRIM29 can promote impairment in repairing double-strand breaks and, consequently, the accumulation of chromosomal instability. Then we studied formation of *TMPRSS2:ERG* fusion genes, a gene fusion that occurs in more than half of PRAD cases, as a model of chromosomal instability (Tomlins et al. 2005). As demonstrated previously, inflammation-related stress (effect of TNFα) promotes accumulation of double-strand DNA breaks in PRAD and accumulation of fusions between the *TMPRSS2* gene and the ETV family of proteins: *TMPRSS2:ERG* and *TMPRSS2:ETV* (Mani et al. 2016). We showed that TRIM29 (LNCaP-TRIM29) overexpression in an LNCaP cell line resulted in a 6-fold decrease in the frequency of *TMPRSS2:ERG* fusions compared to the parental cell line. The effect of TNFα did not lead to considerable growth of fusion rate in LNCaP-TRIM29 but resulted in a 9-fold increase in fusion rate in the LNCaP line (Figure 5A). Since fusion formation is mediated by androgens (Mani et al. 2016), we hypothesized that the addition of TNFα together with testosterone to LNCaP would increase the fusion rate. Indeed, when added with TNFα, testosterone propionate caused a 13-fold increase in gene fusion rate in LNCaP but did not influence fusion rate in LNCaP-TRIM29 (Figure 5A). We also showed that upon overexpression of TRIM29 in LNCaP, the number of γH2AX foci, which are markers of double-strand breaks in DNA, decreases under the effect of TNFα (Anova p-value < 0.05; Figures 5B-D). Altogether, our data support the protective role of TRIM29 from DNA damage under inflammatory stress in studied cell lines.

**Table 1.** TRIM29-interacting proteins in RWPE-1 identified by mass spectrometry. MMR - mismatch excision repair; NER - nucleotide excision repair; Other - other identified genes with known or suspected DNA repair function.

| Gene Name | # of spectral counts |  | # of spectral counts |  | Protein family |
| --- | --- | --- | --- | --- | --- |
|  | IP-TRIM29 |  | IP- IgG |  |  |
| MSH2 | 7 | 15 | 1 | 0 | MMR |
| MSH6 | 6 | 14 | 5 | 3 | MMR |
| RAD23B | 3 | 1 | 0 | 0 | NER |
| CENT2 | 4 | 3 | 0 | 0 | NER |
| DDB1 | 8 | 13 | 3 | 0 | NER |
| CHAF1A | 1 | 3 | 0 | 0 | Other |
| PRPF19 | 15 | 14 | 3 | 3 | Other |
| TP53BP1 | 5 | 2 | 0 | 0 | Other |

**Figure 5.**
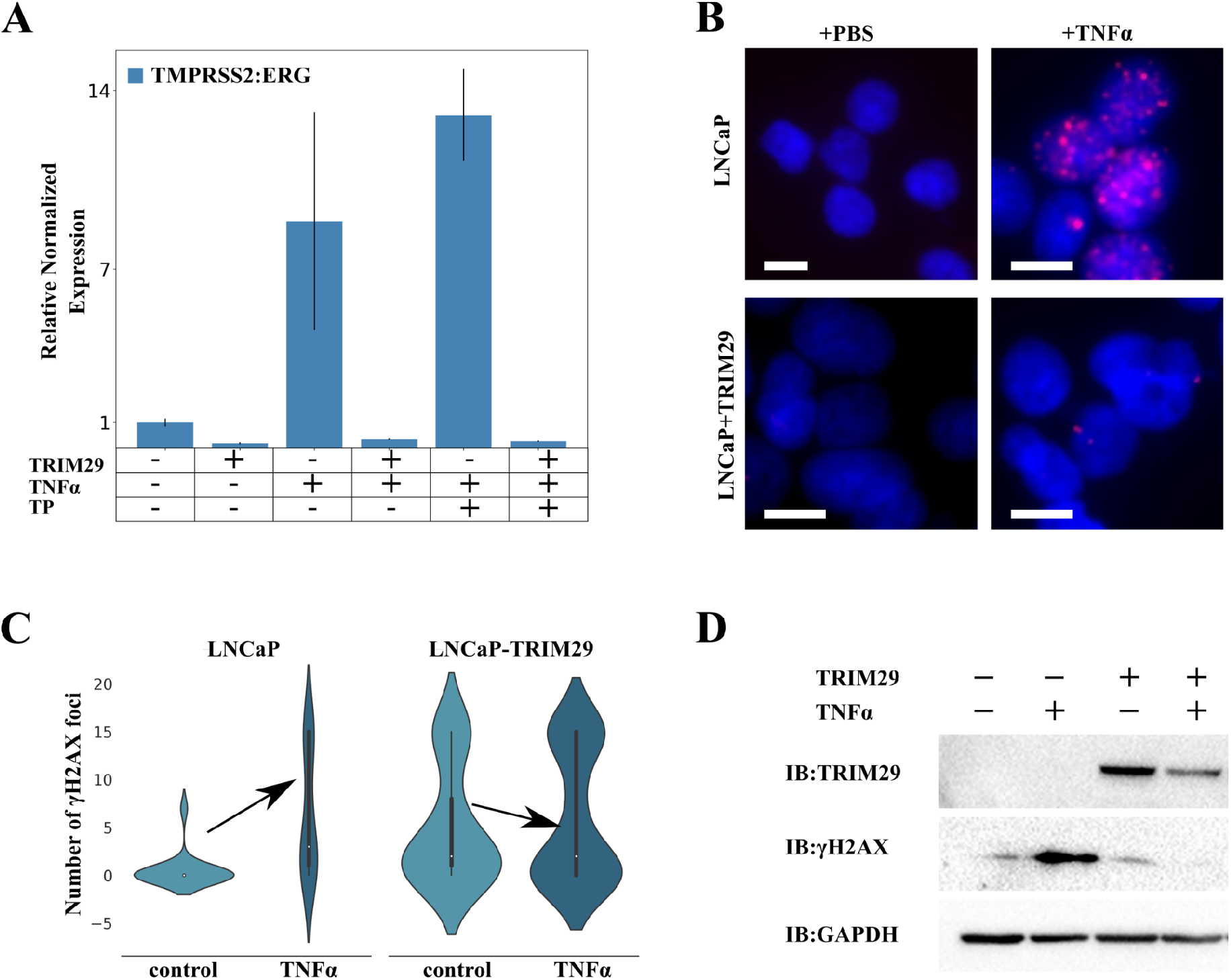
Restoration of TRIM29 expression promotes decrease in genome instability in PRAD. 5A. qRT-PCR analysis of *TMPRSS2:ERG* fusion transcripts from the LNCaP cells with the indicated treatments for 48 h. TP - testosterone propionate. 5B. γH2AX foci formation upon treatment of LNCaP and LNCaP-TRIM29 cells with TNFα (100 ng/mL). Cells were processed 48 h post-treatment. Scale bar indicates 25 μm. 5C. Violin plots of the γH2AX foci number per nucleus in LNCaP and LNCaP-TRIM29 cells before and after TNFα (100 ng/mL) treatment. 5D. Immunoblot (IB) analysis of lysates from LNCaP and LNCaP with stably expressed TRIM29 cells stimulated with TNFα (100 ng/mL).

## Discussion

In this study we performed an integrative analysis of RNA-seq, DNA methylation, and copy number variation data, which revealed a cluster of 143 co-expressed genes associated with epigenomic variability and chromosomal instability. The cluster is enriched with gene signatures of normal basal epithelial cells and is involved in biological processes such as epithelium development and differentiation. We showed that TP63 and TRIM29 regulate the cluster. Transcription factor TP63 is a master regulator of epithelium development, since it regulates the expression of basal markers such as KRT5/6, KRT14, S100A2, and miR-205 (Soares and Zhou 2018). For many squamous-like cancers (pancreas, stomach, head, and neck cancers), TP63 is an oncogene; its high expression is linked with a poor prognosis, while high TP63 expression is associated with a more positive mortality outcome as well as tumor suppressor properties in PRAD (Somerville et al. 2018; Y.-Y. Jiang et al. 2020)(Soares and Zhou 2018; Zhou et al. 2020). TRIM29 (or ATDC) is a ubiquitin ligase of the TRIM family. Similar to TP63, TRIM29 possesses both oncogene and tumor suppressor properties depending on the cancer type (Hatakeyama, 2016). In PRAD TRIM29 expression is considerably lower than normal tissue (Kanno et al. 2014). TRIM29 has been shown to regulate the expression of basal program genes (KRT5, KRT14) through TP63 (Palmbos et al. 2019) in bladder and cervical cancers. We showed that in PRAD, unlike bladder cancer, TP63 binds to the TRIM29 enhancer region but does not regulate TRIM29 expression. Interestingly, knockdown of either TP63 or TRIM29 leads to a decrease in the expression of the TP63 cluster genes. However, overexpression of TRIM29 does not influence the expression of the TP63 gene cluster. But, simultaneous overexpression of TP63 and TRIM29 increases the expression more than TP63 overexpression on its own. Therefore, our study reveals TRIM29 as a modulator of the TP63 transcription program in PRAD. Furthermore, we showed that TP63 and TRIM29 form a complex in the prostate basal epithelium. TRIM29 does not promote TP63 degradation or translocation from the nucleus. Thus, we speculate that TRIM29 owing to its ubiquitin-ligase activity (Q. Li et al. 2018), or susceptible sumoylase activity (Chu and Yang 2011), can post-translationally modify TP63. Previously, a ubiquitin-ligase WWP1 has been demonstrated to influence TP63 transcription factor activity (Peschiaroli et al. 2010). Interestingly, simultaneous expression of TP63 and WWP1 increases the expression of genes KRT14 and KRT5 (Peschiaroli et al. 2010), which concurs with our data from PC3 and bladder cancer cell lines overexpressing TP63 and TRIM29 genes. Moreover, monoubiquitination of the DNA-binding domain of TP53, a TP63 homologue, has been shown to modify TP53 affinity to DNA (Landré et al. 2017). Hypothetically, this can work for TP63 as well due to the high homology of DNA-binding domains of TP63 and TP53. In addition, ubiquitination of TP63 in some cases has been shown to require sumoylation (Ranieri et al. 2018). The sumoylation site is located close to the SAM domain responsible for protein-protein interactions of TP63. Therefore, the sumoylation of TP63 can plausibly affect the choice of protein partners and thus influence its transcription factor activity. Hence, our hypothesis on TRIM29-mediated regulation of TP63-dependent genes is compelling and provides groundwork for further study of TRIM29’s role in post-translational regulation of TP63 activity.

TP63 is a pioneer transcription factor (Bao et al. 2015), which establishes the epigenetic landscape of epidermal enhancers and super-enhancers in squamous-like types of cancer (Somerville et al. 2018; Y.-Y. Jiang et al. 2020; Y. Jiang et al. 2018; Kouwenhoven et al. 2015). Here, for the first time, we demonstrated that TP63 can regulate the level of methylation of enhancers specifically in basal epithelium cells. However, not all the CpG sites associated with the TP63 cluster are subject of demethylation upon expression of TP63 that is supported by previous reports on TP63’s ability to activate enhancers (Somerville et al. 2018; Y.-Y. Jiang et al. 2020). This result can potentially be a consequence of increased hydroxy methylation of cytosines (hmC) at some enhancers by TP63 (Rinaldi et al. 2016). Therefore, further research will be needed to investigate the regulation of hmC levels in enhancers of basal epithelium by TP63. Also, the level of methylation of TP63-dependent CpG sites can be a potential biomarker of TP63 and TRIM29 levels. If so, TP63 and TRIM29 levels can be indirectly evaluated by quantification of CpG methylation in extracellular DNA from urine or blood plasma (Babalyan et al. 2018).

The TP63 cluster is associated with chromosomal instability. Nevertheless, the direct link of the cluster with chromosomal instability is doubtful since the cluster does not contain any genes involved in the DNA repair system. In addition, the cluster genes are not associated with decreased survival upon the effect of DNA-damaging agents (Olivieri et al. 2020; Schleicher et al. 2020) which led us to hypothesize that a single hub gene of the cluster can be associated with chromosomal instability. Prior studies have shown the role of TRIM29 as a scaffold protein of the DNA double-strand break reparation system (Masuda et al. 2015). Notably, DNA reparation system proteins, MSH2/6 and RAD50, are protein partners of TRIM29, which agrees with Masuda’s data for the HeLa cell line (Masuda et al. 2015). Therefore, decreased expression of TRIM29 in PRAD, and perhaps, other cancer types, can cause impairment in DNA repair and consequently increase chromosomal instability. This hypothesis was confirmed in our study using the model of *TMPRSS2:ERG* fusion gene formation in PRAD. In earlier work, the formation of the fusion has been demonstrated to proceed *via* the androgen-dependent pathway and is induced by oxidative or inflammatory stress (Mani et al. 2016). We showed that overexpression of TRIM29 considerably decreases the rates of the *TMPRSS2:ERG* fusion formation even under the effect of inflammatory stress.

Moreover, overexpression of TRIM29 significantly decreased the number of γH2AX foci under the effect of TNFα. As in previous studies, our results show that overexpression of TRIM29 decreased the amount of γH2AX after the effect of ionizing radiation (Masuda et al., 2015). Therefore, our work reveals important insights that link the decreased expression of TRIM29 to increased accumulation of chromosomal instability.

To summarize, we have identified a new partner protein for TP63, TRIM29 ubiquitin ligase, which directly regulates the activity of TP63 as a transcription factor. We have shown TP63- and TRIM29-dependent regulation of basal enhancer methylation and chromosomal instability in PCa. Our findings provide a solid rationale for studying the role of TRIM29 in the regulation of basal epithelium development via TP63 regulation in the prostate gland and epithelium in general.

## Methods

### Data collection

The Cancer Genome Atlas (TCGA) PRAD Due to the high heterogeneity of PRAD data, for the analysis, we selected only samples of patients that meet the following criteria: Gleason scores 6 through 8; no hormonal therapy; over 55 years old (Table S1). Also, we did not include samples marked as contaminated with admixtures of other tissues in TCGA. After quality control and filtration steps, 225 samples of cancerous and 52 normal tissue samples remained.

| Dataset_ID | Biological_source | Treatment | Data_Type | Data_source |
| --- | --- | --- | --- | --- |
| TCGA PRAD | Prostate<br>adenocarcinoma |  | RNA-seq / 450K<br>DNA methylation<br>/ Affymetrix 6.0 | TCGA |
| ICGC PRAD | Prostate<br>adenocarcinoma |  | RNA-seq | ICGC |
| GSE84042 | Prostate<br>adenocarcinoma |  | Affymetrix Human<br>Gene 2.0 ST<br>Array | GEO |
| GSE21032 | Prostate<br>adenocarcinoma |  | Affymetrix Human<br>Exon 1.0 ST<br>Array | GEO |
| GSE26238 | MCF10A | KD TRIM29 | Agilent-014850<br>Whole Human<br>Genome<br>Microarray 4x44K<br>G4112F | GEO |
| GSE71375 | HMEC | KR TRIM29 | Illumina<br>HumanHT-12<br>V4.0 expression<br>beadchip | GEO |
| E-MTAB-1471 | Prostate epithelium | OE TP63, KD<br>TP63 | Agilent Whole<br>Human Genome<br>Microarray 4x44K<br>014850 G4112F | ArrayExpress |
| GSE135257 | ME180 | KD TP63 | RNA-seq | GEO |
| GSE115462 | pancreatic ductal<br>adenocarcinoma | KD TP63, OE<br>TP63 | RNA-seq | GEO |
| GSE115461 | pancreatic ductal<br>adenocarcinoma | TP63,<br>H3K27ac | ChIP-seq | GEO |
| GSE63094 | RWPE-1 | H3K27ac,<br>H3K4me1/3 | ChIP-seq | GEO |
| GSM2400265 | PrEC | DNAseI | DNAseI-seq | GEO |
| GSE57498 | PrEC | H3K27ac,<br>H3K4me1/3 | ChIP-seq | GEO |
| GSE57498 | PC3 | H3K27ac,<br>H3K4me1/3 | ChIP-seq | GEO |
| GSE73783 | LNCaP | H3K27ac,<br>H3K4me1 | ChIP-seq | GEO |
| GSE40050 | LNCaP | H3K4me3 | ChIP-seq | GEO |

### Processing of RNA-seq data

#### TCGA and ICGC data

Level 3 data with mapped readings were used. To build correlation networks, TPM normalization was applied. Only those genes that were expressed in over 50% cancer samples (>1TPM) with expression variability lying above the 25% percentile were included in further analysis. For differential analysis between cancer and not cancer, DESeq2 package was used. Genes were considered differentially expressed, if abs(logFC) > 0.5 and FDR < 0.05.

#### Taylor and Fraser data

Normalized intensities were downloaded with GEO using GEOquery.

Data on knockdown and overexpression of TP63 and TRIM29

#### RNA-seq data

Pre-processing of raw readings was performed with trimmomatic. Then, the readings were mapped onto the genome version hg19 using STAR software. To quantify gene coverage, the featureCounts software with gencode v37 annotation was used. For differential analysis, the DESeq2 package was used. Genes were considered differentially expressed, if abs(logFC) > 0.5 and FDR < 0.05.

#### Hybridization chip data

Normalized intensities were downloaded with GEO using GEOquery. Differential analysis of expression was performed with the Limma package.

### Analysis of DNA methylation data

#### TCGA data

##### The 450K Methylation platform

Files containing raw intensity values (*.idat files) were downloaded from the TCGA portal. To process the data, RnBeads (Assenov et al. 2014) program with the following setup parameters was used: methylumi.noob (Davis et al. 2014) + BMIQ (Teschendorff et al. 2013) normalization; limma algorithm for differential analysis. CpG site was considered differentially methylated if abs(deltaBeta)>0.2 and FDR<0.05.

#### Data on TP63 overexpression in PC3

##### EPIC Methylation platform

To process the data, RnBeads (Assenov et al. 2014) program was used with the following setup parameters: ENmix (Xu et al. 2016) + BMIQ (Teschendorff et al. 2013); limma algorithm for differential analysis (Ritchie et al. 2015). To search for motifs of transcription factors, HOMERv4 (Heinz et al. 2010) software with the motif database HOCOMOCOv11 (Kulakovskiy et al. 2018) were used.

##### Analysis of CNV data

Level 3 open data obtained on Affymetrix 6.0 platform were used. A region was considered to have an altered copy number if the absolute value of the ratio of intensities between sample and control exceeded 0.3. To describe chromosomal instability, chromosomal instability index (CIN), equal to the ratio of total length of CNV to the length of a chromosome, averaged across all chromosomes, was used.

### Determination of clusters of co-expressed genes associated with epigenomic variability and chromosomal instability

To generate clusters, we used the WGCNA package (Langfelder and Horvath 2008) with standard parameters. We found 13 clusters of co-expressed genes in TCGA data. For each cluster, we determined its expression pattern, cluster eigengene (EG), which is the main component of PCA.

As it has been demonstrated in a previous work, the level of methylation of CpG sites can strongly depend on the ratio of various types of prostate cells in a sample (Pellacani et al. 2018). We used the MuSig package (X. Wang et al., 2019) and prostate scRNA-seq data (Henry et al. 2018) to calculate the ratio of stromal, basal, and luminal epithelium cells in prostate gland in TCGA samples. We used calculated ratios as co-variants for further regression analysis. To determine the set of non-overlapping CpG sites associated with each cluster, we used the following regression model:

EGi = ai + bi*CpGj + Covi + Erri,

where EGi is cluster i eigengene; ai, a constant; bi, coefficient by the vector of methylation of CpGi; and Cov, co-variants that include the ratio between various types of cells in a sample and the rest eigengenes of the cluster except for the studied one to determine only CpG sites uniquely associated with the cluster. We thus determined 14443 differentially methylated CpG sites associated with eigengenes of the clusters.

We used linear regression to determine gene clusters associated with CIN:

CIN = ai + bi*EGi + Covi + Erri,

where CIN is chromosomal instability index, EGi, is cluster i eigengene; ai, a constant; bi, a coefficient at EGi; Cov, co-variants that include the ratio between various types of cells in a sample.

To determine association of clusters with differentially expressed genes between cancer and normal tissue, we used GSEA, where genes of each cluster were used as signatures.

#### Super-enhancer-associated genes prediction

We used the ROSE framework (Hnisz, Day, and Young 2016) with DNAse-seq and H3K27ac ChIP-seq for normal prostate epithelial cell line PrEC for SEs prediction. For the prediction link between genes and enhancers and SEs, we used the Activity-by-contact framework (Fulco et al. 2019).

#### Cell culture

Cells were cultured in corresponding media in a 5% CO_2_ incubator at 37°C as described (paper cite). The cells were washed with Hank’s solution to remove dead cells and the medium was replaced with fresh one every 3 days.

#### Preparation of lentivirus particles

One day prior to transfection, Phoenix cells were inoculated onto 10-cm Petri dishes covered with 0.1% gelatin 7 × 10^5^ cells per dish. The cells were transfected with auxiliary plasmids containing the Rev (15.3% by mass to total DNA), RRE (29.8% by mass to total DNA), and VSV-F (5.6% by mass to total DNA) genes and target plasmid encoding TRIM29_FLAG. Transfection was performed using the TurboFect (ThermoFischer Scientific, US) transfection agent, 26.4 µL per 13.2 µg DNA. The procedure of transfection was performed in accordance with the manufacturer’s recommendations. Virus-containing supernatant was collected 24, 48, and 72 h after transfection, filtered through 0.45 µm filter and stored at –70°С.

#### Cell line generation

Two days before transduction, cells were seeded into 24-well plate 2 × 10^4^ cell per well. One hour prior to transduction, 8 µg/mL polybrene was added. Cells were transduced with virus particles containing TP63-FLAG or TRIM29-FLAG. After 24 h, cell medium was replaced. Two days after transfection, cells were trypsinized and TP63/TRIM29-infected cells were isolated on a FACS BD Aria III cell sorter.

#### Cell treatment with TNFα

Prior to treatment with TNFα, cells were cultured to 50–60% confluence layer. TNFα was introduced into the medium to a final concentration of 100 ng/mL and cultured in an incubator for 48 hours.

#### Chromatin immunoprecipitation

Chromatin immunoprecipitation was performed using the SimpleChIP Plus Enzymatic Chromatin IP Kit (Cell Signaling) according to the manufacturer’s manual. In brief, 4 × 10^6^ cells were fixed with formaldehyde, cell nuclei were isolated, and chromatin was fragmented with micrococcal nuclease. Then, chromatin was incubated with 5 µL anti-TP63 antibodies (Cell Signaling #13109) overnight and TP63-bound DNA was precipitated on magnetic beads. The DNA was purified from bound protein complexes and used for massive parallel sequencing using the Illumina HiSeq-2500 or for quantitative PCR.

The following algorithm was used to analyze chromatin immunoprecipitation followed by sequencing: pre-processing of raw reads was performed by trimmomatic tool; then the reads were mapped onto human genome version hg19 using the bwa mem program with standard parameters. The *.bam files thus obtained were sorted and indexed with the samtools program. Reliably determined regions of TP63 or histone binding with determined modifications were found using the macs2 program. ChIP-seq data were visualized using the deepTools program.

Paired-end libraries were prepared according to the manufacturer’s recommendations using NEBNext Ultra II DNA Library Prep Kit (New England Biolabs, USA). The libraries were indexed with NEBNext Multiplex Oligos kit for Illumina (96 Index Primers, New England Biolabs, USA). Size distribution for the libraries and their quality were assessed using a high-sensitivity DNA chip (Agilent Technologies). The libraries were subsequently quantified by Quant-iT DNA Assay Kit, High Sensitivity (Thermo Scientific, USA). DNA sequencing was performed on the HiSeq 2500 platform (Illumina, USA) according to the manufacturer’s recommendations, using the following reagent kits: HiSeq Rapid PE Cluster Kit v2, HiSeq Rapid SBS Kit v2 (200 cycles), HiSeq Rapid PE FlowCell v2 and a 1% PhiX spike-in control.

#### TP63 and TRIM29 knockdown

TP63 and TRIM29 knockdown was performed using RNA interference. siRNA was transfected into RWPE-1 cells in a 24-well plate using HiPerFect Transfection Reagent (Qiagen, 301704). siRNA, 68 ng, was mixed with 2 µL transfection reagent in 100 µL of Opti-MEM™ (Gibco) medium; the mixture was incubated at room temperature for 10 min. RWPE-1 cells were seeded onto a 24-well plate at 3.5 × 10^4^ cells/cm^2^. Cells were introduced into wells in 400 µL Keratinocyte SFM (Gibco) medium and the mixture of siRNA and transfection reagent was added to the cells immediately. Therefore, total medium volume was 500 µL in a well; final siRNA concentration, 10 nM. After 24 hours, the medium was partially replaced with fresh Keratinocyte SFM (Gibco). The efficiency of transfection was evaluated 48 h after the addition of siRNA by western blotting.

#### Cell lysate preparation for western blot analysis

Cell cultures were collected, and 1 × 10^6^ cells were counted by trypan blue exclusion and washed with ice cold PBS. Cells were then resuspended in 100 μL PBS and lysed with 100 μL of 2x Laemmli Sample Buffer supplemented with β-mercaptoethanol by boiling for 5-10 min.

#### Isolation of cytoplasm and nuclear fractions

Cells were washed with PBS and removed with a scraper in cool buffer A (0.35 М sucrose, 2 мМ MgCl_2_, 0.1 mM EDTA, 0.1% Triton X-100, 10 mM Tris-HCl pH 8.0, 1 mM DTT, 0.1 mM PMSF, and protease inhibitors). Cell lysate was passed 10 times through a 21G needle syringe and a 23G syringe, 3 times. Then, it was centrifuged at 2000 g for 10 min. The supernatant was collected as a cytoplasm fraction.

The precipitate containing nuclei was washed with 0.5 M sucrose and centrifuged at 10000 g for 10 min. The precipitate was resuspended in buffer C (400 mM NaCl, 20 mM HEPES pH 7.9, 25% glycerol, 1.5 mM MgCl_2_, 0.4 mM EDTA, 0.4 mM PMSF, 1 mM DTT, protease inhibitors) and passed 5 times through a 21G needle syringe. Then, it was incubated at +4°С for 40 min and centrifuged at 12000 g for 30 min. The supernatant was collected as the nuclear fraction.

#### Immunoprecipitation

Cells were washed with PBS solution. Cell lysis buffer (Cell Signaling Technology, Great Britain) was added and incubated for 5 min on ice. Cells were removed with a scraper. Cell lysate was homogenized on an ultrasound bath. Lysate was centrifuged at 10000 g for 10 min. The supernatant was incubated with antibodies overnight at +4°С while stirring. Lysate with antibodies was incubated with magnetic beads for 1 h while stirring at room temperature. Magnetic beads were washed 5 times with cell lysis buffer.

For western blotting, the proteins were eluted with Laemmli buffer at 95°С for 5 min. Total protein concentration in the sample was determined using Bradford assay.

To prepare samples for mass spectrometry analysis, proteins were eluted with elution buffer (8 М urea, 2 М thiourea, 10 mM Tris/HCl pH 8.0) on a shaker at 25°С for 2 h.

#### Western blotting

Electrophoretic separation of proteins was carried out in a Bio-Rad Mini Protean chamber. Then, semidry electrotransfer of proteins to a PVDF membrane was carried out in a Trans-Blot Turbo Transfer System (Bio-Rad, USA). The membrane was incubated in blocking buffer (PBST, 5% non-fat dry milk). Then, it was incubated with primary antibodies for 1 h at room temperature or at +4°С overnight, washed in PBST solution, and incubated with a solution of secondary antibodies conjugated with horseradish peroxidase for 1 h at room temperature or at +4°С overnight. Proteins were visualized using Pierce ECL Western Blotting Substrate (Thermo Fisher Scientific, USA) at Chemidoc (Biorad, USA).

#### Sample preparation for mass spectrometric analysis

After immunoprecipitation, DTT was added to the eluted samples to a final concentration of 5 mM and incubated for 30 min on a shaker at room temperature to restore disulfide bonds. Freshly prepared iodoacetamide was added to a final concentration of 10 mM and incubated in the dark for 20 min to alkylate the thiol groups of cysteine. The mixture was diluted with 35 mM ammonium carbonate solution 4-fold. Sequencing Grade Modified Trypsin (Promega, USA) was added 0.1 μg trypsin per 10 μg protein and incubated on a shaker at 37°С overnight. Trypsin was neutralized with a 5-fold volume of a 5% formic acid solution. Desalting was performed using reversed phase chromatography on homemade StageTips with SDB-RPS filter according to the protocol from (Rappsilber et al., 2003). The samples were concentrated in a vacuum centrifuge and redissolved in 3% acetonitrile with 0.1% trifluoroacetic acid solution. The amount of protein in the sample was approximately 10 μg.

#### LC-MS analysis of tryptic peptides

After trypsinolysis, peptide fractions were loaded onto a column (diameter 75 μm, length 50 cm) with an Aeris Peptide XB-C18 2.6 μm sorbent (Phenomenex) in an aqueous solution containing 3% acetonitrile and 0.1% trifluoroacetic acid. Separation of peptides was performed on an Ultimate 3000 Nano LC System (Thermo Fisher Scientific), coupled to a Q Exactive HF mass spectrometer (Thermo Fisher Scientific) using a nanoelectrospray source (Thermo Fisher Scientific). Peptides were loaded onto a heated 40°C column in buffer A (0.2% formic acid (FA) in water) and eluted with a linear (120 min) gradient 4 --> 55% buffer B (0.1% FA, 19.9% water, 80% acetonitrile) in A at a flow rate of 350 nL/min. Before each new load, the column was washed with 95% buffer B in A for 5 min and equilibrated with buffer A for 5 min.

Mass spectrometry data were saved with automatic switching between MS1 scans and up to 15 MS/MS scans (topN method). The target value for the MS1 scan was set to 3 × 10^6^ in the range of 300−1200 *m*/*z* with a maximum ion injection time of 60 ms and a resolution of 60000. The precursor ions were isolated with a window width of 1.4 *m*/*z* and a fixed first mass of 100,0 *m*/*z*. The precursor ions were fragmented by high-energy dissociation in a C-trap with a normalized collision energy of 28 eV. MS/MS scans were saved with a resolution of 15000 at *m*/*z* 400 and at a value of 1 × 10^5^ for target ions in the range of 200−2000 *m*/*z* with a maximum ion injection time of 30 ms.

#### Analysis of LC-MS data

The conversion of the “raw” mass spectrometric data from the instrument into MGF (Mascot Generic Format) mass sheets was carried out using the ProteoWizard msconvert utility with the following parameters: MS Levels 2-2, Peak Picking 2-2, Threshold Peak Filter Absolute intensity - Most intense - 1, Zero Samples 2-2.

For identification and quantitative analysis of protein partners, the MaxQuant program (v1.5.3.30) was used with the Andromeda algorithm against the protein database UniProt Knowledgebase (UniProtKB), the human taxon, with the following parameters: the accuracy of determining the parent and daughter ions was 20 and 50 ppm, respectively; protease, trypsin; one missed cleavage per peptide is possible; variable modification of methionine, oxidation; fixed modification of cysteine, carbamidomethylation. The reliability of identification of both peptides and proteins was limited to 1% FDR, which was determined using the “target decoy” approach. For quantitative label-free analysis, LFQ values were calculated using the MaxQuant software.

#### Immunocytochemical analysis

Cells were washed with PBS solution 2 times for 5 min and fixed with 4% PFA for 30 min. The cells were washed 2 times for 5 min. Membranes were permeabilized with 0.1% Triton X-100 in PBS for 5 min. Nonspecific antigen adsorption was blocked by washing in 0.1% Tween 20 PBS solution 3 times for 5 min and then in a block solution (PBS, 0.1% Tween 20, 5% FBS, 5% goat serum) for 30 min. A solution of primary antibodies was added in a block solution and incubated for 2 h at room temperature. The cells were washed from primary antibodies with a solution of 0.1% Tween 20 3 times for 5 min. A solution of secondary antibodies labeled with the Alexa Fluor 555 fluorophore was added in PBS and incubated in the dark for 1 h at room temperature.

Then, the cells were washed from secondary antibodies with a solution of 0.1% Tween 20 3 times for 5 min. To stain the nuclei, the mixture was incubated with DAPI 100 ng/mL for 10 min in the dark at room temperature and then washed with PBS solution once. PBS was removed and glycerol was applied to the sample prior to covering with a coverslip. The stained proteins were visualized using a fluorescence microscope (Nikon Eclipse Ni-E, Japan).

#### DNA isolation and bisulfite conversion

Cells (∼10^6^) were resuspended in a lysis buffer (10mM Tris pH=8.0, NaCl 100mM, 10mM EDTA pH=8.0, 0.5% SDS) and incubated with Proteinase K overnight. For DNA purification the phenol chloroform extraction was used. The DNA was then bisulfite converted using the EpiMark® Bisulfite Conversion Kit (E3318S, NEB) according to the manufacturer’s protocol.

#### Infinium Methylation EPIC Beadchip array

All DNA methylation experiments were performed according to Illumina manufacturer instructions for the Infinium Methylation EPIC 850K BeadChip Array (Illumina, USA). EPIC BeadChips were imaged using the Illumina iScan System (Illumina, United States).

#### Isolation of RNA and cDNA synthesis

Isolation of RNA from cells was performed using the Lira reagent (Biolabmix, Russia) according to the manufacturer’s protocol. The quality of the isolated RNA was assessed using agarose gel electrophoresis. The concentration of the resulting RNA preparation was measured spectrophotometrically on an Infinite 200 Pro M Plex plate reader (Tecan, Switzerland) determining the absorbance at 260 nm. The purity of the preparation was assessed by the ratio of absorption at wavelengths of 260 and 230 nm.

A mixture was prepared containing 1 μg of RNA, 1 U DNase I (Thermo Fisher Scientific, USA), and a reaction buffer with MgCl_2_. Incubation was carried out for 30 min at 37°C. Then, 1 μl of 50 μM EDTA was added and the solution was heated to 65°C to inactivate DNase.

The first strand of cDNA was synthesized from a single-stranded RNA template using the MMLV RT kit (Evrogen, Russia) according to the manufacturer’s protocol.

#### RNA Sequencing

Library preparation was performed with NEBNext Poly(A) mRNA Magnetic Isolation Module and NEBNext Ultra II Directional RNA Library Prep Kit (NEB) according to the manufacturer’s protocol. The library underwent a final cleanup using the Agencourt AMPure XP system (Beckman Coulter) after which the libraries’ size distribution and quality were assessed using a high sensitivity DNA chip (Agilent Technologies). Libraries were subsequently quantified by Quant-iT DNA Assay Kit, High Sensitivity (ThermoFisher). Finally, libraries were sequenced by a high throughput run on the Illumina HiSeq 2500 using 2 × 125 bp paired-end reads.

#### Real-time PCR

Real-time PCR was performed in a volume of 20 μL using the obtained cDNA as a template, qPCRmix-HS SYBR (Evrogen, Russia) containing HS Taq DNA polymerase, SYBR Green I dye, a mixture of deoxynucleoside triphosphates, Mg^2+^, reaction buffer, and primers for the target gene (TMPRSS2_ERG_fwd and TMPRSS2_ERG_rev) and a reference gene (ACTB_fwd and ACTB_rev). Each sample was prepared four times.

#### MSRE-PCR

The protocol was used from the article (Melnikov et al. 2005) with minor modifications. HpaII was used as a methyl-sensitive restriction enzyme. 500ng genomic DNA was incubated with a restriction enzyme overnight. The treated DNA was purified by phenol-chloroform extraction.

The purified DNA was used as a template for quantitative analysis. The region of the GAPDH gene in which there are no HpaII restriction sites was used as a control.

#### Statistics or Data analysis

Results were analyzed using 2-tailed Student’s t-test, Fisher’s exact test and 2-way ANOVA. P-value less than 0.05 was considered statistically significant.

## Data and Software Availability

The accession number for the ChIP-seq, RNA-seq and DNA methylation data reported in this paper is GEO: GSE204813

## Acknowledgements

The research was supported by grant 075-15-2019-1669 from the Ministry of Science and Higher Education of Russian Federation; part of this work was supported by the Russian Foundation for Basic Research projects [17-29-06063].

## Author contributions

RS and GA conceived the study and RS designed all experiments and analyzed the data. OZ and AM performed most experiments. AF generated expression vectors. AB performed ChIP-seq experiments. KS and AL performed DNA and RNA sequencing and DNA-methylation analysis. TP and TZ generated siRNA oligos. RS, GA, ML, EG, ES and VG interpreted the results. RS wrote the manuscript. GA and ML supervised the project.

## Supplementary figures and tables

Table S1 related to Figure 1

Data sets used in the study

Sheet1- List of all datasets used in the study

Sheet2 - List of PRAD sample IDs used in the study

Table S2 related to Figure 2

TP63 regulates the gene cluster associated with epigenetic variability and chromosomal instability in PRAD

Sheet1 - list of genes from TP63 cluster

Sheet2 - Ontology analysis of the TP63 cluster genes. Gene Ontology (GO) terms related to biological process

Sheet3 - Ontology analysis of the TP63 cluster genes. Gene Ontology (GO) terms related to cellular components

Table S3 related to Figure 3

CpG sites associated with TP63 cluster belong to TP63-dependent enhancers and super-enhancer

Sheet1 - list of CpG sites associated with TP63 cluster

Sheet2 - Ontology analysis of genes located near the TP63 CpG sites. Gene Ontology (GO) terms related to biological process

Sheet3 - Ontology analysis of genes located near the TP63 CpG sites. Disease ontology

Table S4 related to Figure 4

TRIM29 interacts with TP63 and regulates expression of the TP63 cluster

Sheet1 - Ontology analysis of the TP63 cluster genes under TP63 and TRIM29 simultaneously regulation. Gene Ontology (GO) terms related to biological process

Table S5 related to Figure 5

TRIM29 promotes decrease of chromosomal instability in PRAD

Sheet1 - Partners of TRIM29 in RWPE-1 cells

Sheet2 - Association between the TP63 cluster and the response to genotoxic stress

**Figure S1 related to Figure 2.**
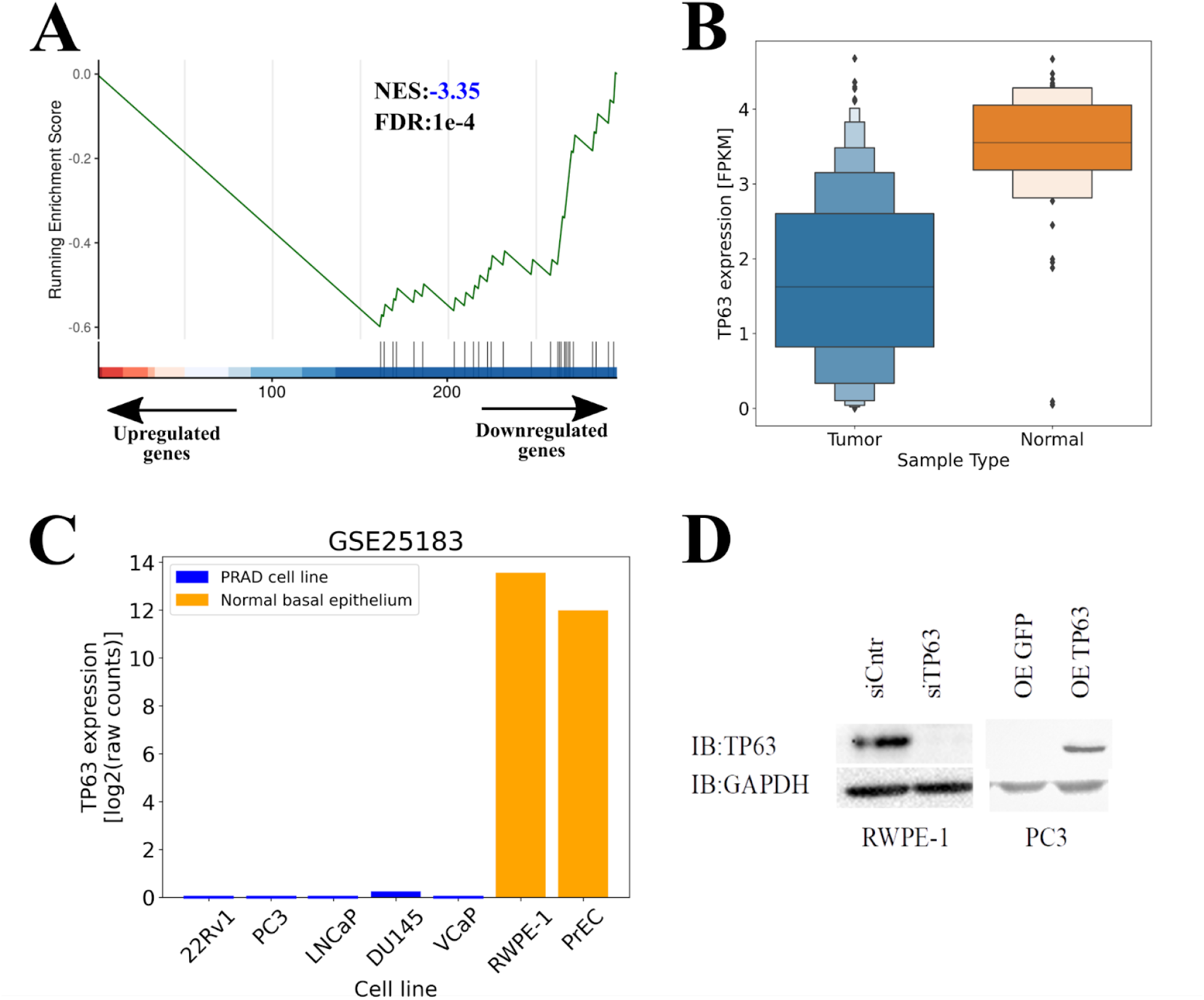
A. GSEA plots evaluating the TP63 cluster signatures upon differential expressing genes between cancer and normal samples in TCGA dataset. B. Boxenplot represents expression level of *TP63* in TCGA PRAD samples. Each successive level outward 50%-percentile contains half of the remaining data. C. Expression of *TP63* in PRAD and normal prostate epithelium cell lines. D. Immunoblot (IB) analysis of overexpression (OE) of *TP63* in PC3 cells and knockdown (KD) of *TP63* in the RWPE-1 cells.

**Figure S2 related to Figure 3.**
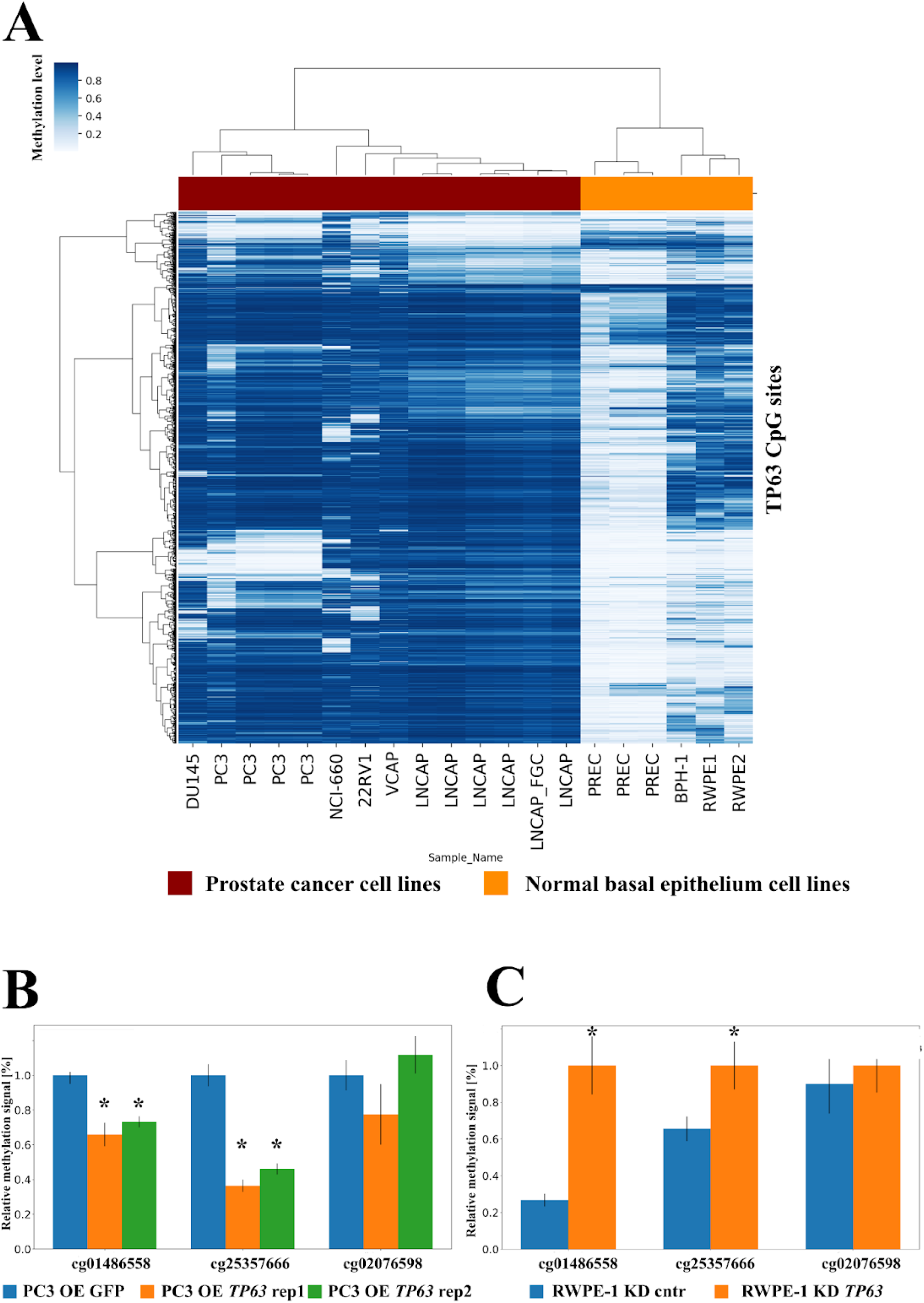
A. Representation of DNA methylation level of 1645 TP63 CpG sites in normal and PRAD cell lines. B. Overexpression of *TP63* in PC3 promotes decreased methylation of two CpG sites (MRSE-qPCR) compared to an empty vector control. Error bar = SD. N = 3 C. *TP63* knockdown in the RWPE-1 cell line promotes increased methylation of two CpG-sites (MRSE-qPCR) compared to control. Error Bar = SD. N = 3

**Figure S3 related to Figure 4.**
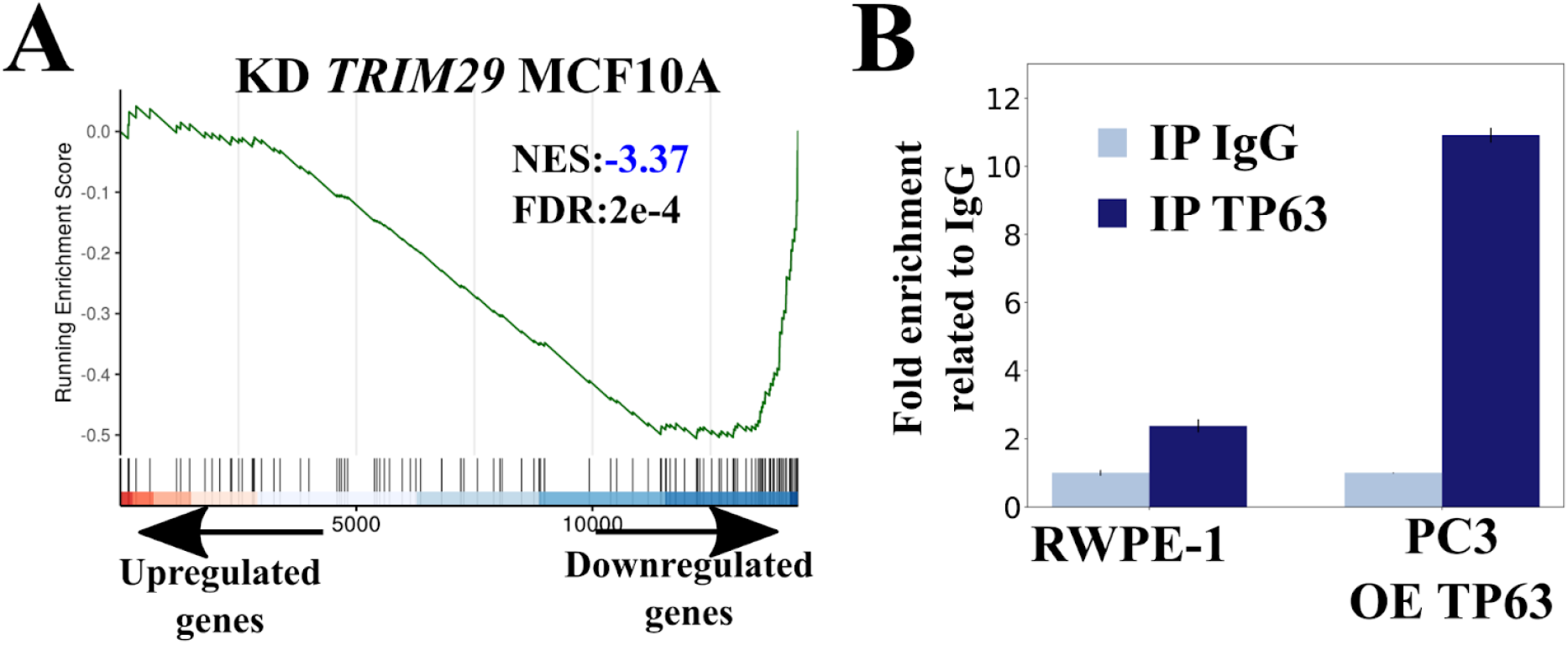
A - GSEA plots evaluating the TP63 cluster signature upon *TRIM29* knockdown in MCF10A (breast basal epithelium, GSE26238). B - TP63 ChIP in RWPE-1 and PC3 cells with *TP63* overexpression demonstrates enrichment of *TRIM29* enhancer 1. All samples were done in triplicate. Error Bars = SEM IP - immunoprecipitation.

